# Regulation by cyclic-di-GMP attenuates dynamics and enhances robustness of bimodal curli gene activation in *Escherichia coli*

**DOI:** 10.1101/2022.05.23.493020

**Authors:** Olga Lamprecht, Maryia Ratnikava, Paulina Jacek, Eugen Kaganovitch, Nina Buettner, Kirstin Fritz, Ina Biazruchka, Robin Köhler, Julian Pietsch, Victor Sourjik

## Abstract

Curli amyloid fibers are a major constituent of the extracellular biofilm matrix formed by bacteria of the Enterobacteriaceae family. Within *Escherichia coli* biofilms, curli gene expression is limited to a subpopulation of bacteria, leading to heterogeneity of extracellular matrix synthesis. Here we show that bimodal activation of curli expression occurs not only in submerged and macrocolony biofilms, but also in well-mixed planktonic cultures of *E. coli*, resulting in all-or-none stochastic differentiation into distinct subpopulations of curli-positive and curli-negative cells at the entry into the stationary phase of growth. Stochastic curli activation in individual *E. coli* cells could further be observed during continuous growth in a conditioned medium in a microfluidic device, which further revealed that the curli-positive state is only metastable. In agreement with previous reports, regulation of curli gene expression by c-di-GMP via two pairs of diguanylate cyclase and phosphodiesterase enzymes, DgcE/PdeH and DgcM/PdeR, modulates the fraction of curli-positive cells under all tested growth conditions. Unexpectedly, removal of this regulatory network does not abolish the bimodality of curli gene expression, although it affects dynamics of activation and increases heterogeneity of expression levels among individual cells. Moreover, the fraction of curli-positive cells within an *E. coli* population shows stronger dependence on growth conditions in the absence of c-di-GMP regulation. We thus conclude that, while not required for the emergence of bimodal curli gene expression in *E. coli*, this c-di-GMP regulatory network attenuates the frequency and dynamics of gene activation and increases its robustness to cellular heterogeneity and environmental variation.

## Introduction

Curli amyloid fibers are the key component of the extracellular matrix produced during biofilm formation by *Escherichia coli*, *Salmonella enterica*, and other Enterobacteriaceae [1–9]. In *E. coli* and *S. enterica* serovar Typhimurium, curli genes are organized in two divergently transcribed *csgBAC* and *csgDEFG* operons that share a common intergenic regulatory region [10]. Expression of these operons is under regulation of the stationary phase sigma factor σ^S^ (RpoS) and thus becomes activated during the entry into the stationary phase of growth [4, 11–14]. This activation is achieved by the σ^S^-dependent induction of the transcriptional regulator CsgD, which then controls the expression of the *csgBAC* operon that encodes the major curli subunit CsgA along with the curli nucleator CsgB and the chaperone CsgC [7, 8, 15]. In turn, *csgD* expression in *E. coli* and *S.* Typhimurium is either directly or indirectly regulated by multiple cellular factors that mediate responses to diverse environmental changes, including both global and specific transcriptional regulators, small regulatory RNAs and second messengers (reviewed in [16–19]).

One of the key regulators of *csgD* is the transcription factor MlrA [13, 14, 20, 21]. The activity of MlrA depends on cellular levels of bacterial second messenger bis-(3’-5’)-cyclic dimeric guanosine monophosphate (c-di-GMP), and in *E. coli* this control is known to be mediated by a pair of the interacting diguanylate cyclase (DGC) and phosphodiesterase (PDE) enzymes, DgcM and PdeR, that form a ternary complex with MlrA [12, 14, 22]. MlrA is kept inactive by binding PdeR, and this interaction is relieved when the latter becomes active as a PDE thus acting as the trigger enzyme [22, 23]. This inhibition is counteracted both by DgcM, which locally produces c-di-GMP to engage PdeR, as well as by the global pool of c-di-GMP. Besides its enzymatic activity, DgcM might also activate MlrA through direct protein interaction. Another DGC-PDH pair, DgcE and PdeH, provides global regulatory input into the local DgcM-PdeR-MlrA regulation [12, 24]. At least under typical conditions used to study *E. coli* biofilms, this regulatory network appears to be the only c-di-GMP-dependent input that controls *csgD* expression [24].

Previous studies of *E. coli* macrocolony biofilms formed on agar plates showed that curli expression occurs in the upper layer of the colony, but even in this layer its expression remained heterogeneous [25–27], indicating an interplay between global regulation of curli gene expression by microenvironmental gradients within biofilms and its inherent stochasticity. Differentiation of *E. coli* into distinct subpopulations of cells either expressing or not expressing curli was also observed in submerged biofilms formed in liquid cultures, whereby curli expression was associated with cellular aggregation [28]. Furthermore, bi- or multimodality of *csgD* reporter activity was also observed in the early stationary phase among planktonic cells in *S.* Typhimurium [29, 30] and *E. coli* [27]. Given established c-di-GMP-dependent regulation of CsgD activity, it was proposed that bistable curli expression originates from a toggle switch created by mutual inhibition between DgcM and PdeR, which could act as a bistable switch on *csgD* gene expression [27, 31].

In this study we demonstrate that differentiation of *E. coli csgBAC* operon expression, leading to formation of distinct subpopulations of curli-positive and -negative cells, occurs not only in submerged and macrocolony biofilms but also stochastically in a well-stirred planktonic culture and thus in the absence of any environmental gradients. Similar stochastic and reversible differentiation could be further observed among cells growing in conditioned medium in the microfluidic device. Although the c-di-GMP regulatory network consisting of the DgcE/PdeH and DgcM/PdeR pairs modulates the fraction of curli-positive cells in the population and the dynamics and uniformity of curli gene activation, it is not required to establish the bimodality of their expression.

## Materials and methods

### Bacterial strains and plasmids

All strains and plasmids used in this study are listed in Table S1. A derivative of *E. coli* W3110 [26] that was engineered to encode a chromosomal transcriptional sfGFP reporter downstream of the *csgA* gene [28] (VS1146) was used here as the wild-type strain. Gene deletions were obtained with the help of P1 phage transduction using strains of the Keio collection [32] as donors, and the kanamycin resistance cassette was removed using FLP recombinase [33]. For expression, *dgcE* and *pdeH* genes were cloned into the pTrc99A vector [34].

### Growth conditions for planktonic cultures

Planktonic *E. coli* cultures were grown in tryptone broth (TB) medium (10 g tryptone, 5 g NaCl per liter, pH 7.0) or lysogeny broth (LB) medium without salt (10 g tryptone, 5 g yeast extract, per liter, pH 7.0), supplemented with antibiotics where necessary. Overnight cultures were inoculated from LB agar plates, grown at 30°C and diluted 1:100, unless indicated otherwise, in 5-10 ml of fresh TB and grown at 30°C at 200 rpm in 100 ml flasks in a rotary shaker until the indicated OD_600_ or overnight (18-25 h; OD_600_ ~ 1.3-1.8). Alternatively, cultures were grown in TB in 96-well plates with orbital shaking in a plate reader, with 200 μl culture per well. Where indicated, bacterial cultures were supplemented with either 1 mM L-serine (after 6 h of growth) or 0.1–10 mg/l DL-serine hydroxamate at inoculation.

### Growth and quantification of submerged biofilms

Submerged biofilms were grown and quantified as described previously [28], with minor modifications. Overnight bacterial cultures grown in TB were diluted 1:100 in fresh TB medium and grown at 200 rpm and 30°C in a rotary shaker to OD_600_ of 0.5. The cultures were then diluted in fresh TB medium to a final OD_600_ of 0.05, and 300 μl was loaded into a 96-well plate (Corning Costar, flat bottom; Sigma-Aldrich, Germany) and incubated without shaking at 30°C for 46 h.

For quantification of biofilm formation, the non-attached cells were removed and the wells were washed once with phosphate-buffered saline (PBS; 8 g NaCl, 0.2 g KCl, 1.44 g Na_2_HPO_4_, 0.24 g KH_2_PO_4_). Attached cells were fixed for 20 min with 300 μl of 96% ethanol, allowed to dry for 40 min, and stained with 300 μl of 0.1% crystal violet (CV) solution for 15 min at room temperature. The wells were subsequently washed twice with 1x PBS, incubated with 300 μl of 96% ethanol for 35 min and the CV absorption was measured at OD_595_ using INFINITE M NANO^+^ plate reader (Tecan Group Ltd., Switzerland). Obtained CV values were normalized to the OD_600_ values of the respective biofilm cultures.

### Macrocolony biofilm assay

Macrocolony biofilms were grown as described previously [26]. Briefly, 5 μl of the overnight liquid culture grown at 37°C in LB medium (10 g tryptone, 5 g NaCl, and 5 g yeast extract per liter) was spotted on salt-free LB agar plates supplemented with Congo red (40 μg/ml). Plates were incubated for 8 days at 28°C.

### Fluorescence measurements

Measurements of GFP expression in an INFINITE M1000 PRO plate reader (Tecan Group Ltd., Switzerland) were done using fluorescence excitation at 483 nm and emission at 535 nm. Relative fluorescence was calculated by normalizing to corresponding OD_600_ values of the culture.

For fluorescence measurements using flow cytometry, aliquots of 40-300 μl of liquid bacterial cultures were mixed with 2 ml of tethering buffer (10 mM KH_2_PO_4_, 10 mM K_2_HPO_4_, 0.1 mM EDTA, 1 μM L-methionine, 10 mM lactic acid, pH 7.0). Macrocolonies were collected from the plate, resuspended in 10 ml of tethering buffer and then aliquots of 40 μl were mixed with 2 ml of fresh tethering buffer. Samples were vigorously vortexed and then immediately subjected to flow cytometric analysis using BD LSRFortessa Sorp cell analyzer (BD Biosciences, Germany) using 488-nm laser. In each experimental run, 50,000 individual cells were analyzed. Absence of cell aggregation was confirmed by using forward scatter (FSC) and side scatter (SSC) parameters. Data were analyzed using FlowJo software version v10.7.1 (FlowJo LLC, Ashland, OR, US).

### Microfluidics

Conditioned medium was prepared by cultivating wild-type *E. coli* in TB in a rotary shaker at 30°C for 20 h, after which the cell suspension was centrifuged at 4000 rpm for 10 min, medium was filter-sterilized and stored at 4°C. Mother machine [35] microfluidics devices were designed, fabricated and operated as described in Supporting protocols. *E. coli* cells from the overnight culture in TB were loaded into the mother machine by manual infusion of the cell suspension through one of the two inlets using a 1-ml syringe. Cells were first allowed to grow at 30 °C for 4 h in fresh TB, then switched to the conditioned TB and cultivated for up to 26 h. Phase contrast and GFP fluorescence images were acquired using a Nikon Eclipse Ti-E inverted microscope with a time interval of 10 min. Details of image analysis are described in Supporting protocols.

## Results

### Bimodal curli gene expression is induced in planktonic culture

In order to characterize curli expression in planktonic culture of *E. coli*, we followed the induction of a chromosomal transcriptional reporter for the *csgBAC* operon, where the gene encoding for stable green fluorescent protein (GFP) was cloned downstream of *csgA* with a strong ribosome binding site as part of the same polycistronic RNA [28]. In our previous study of submerged *E. coli* biofilms, this reporter showed bimodal expression both in the surface-attached biofilm and in the pellicle at the liquid-air interface, and its activity correlated with the recruitment of individual cells into multicellular aggregates [28]. When *E. coli* culture was grown at 30°C in tryptone broth (TB) liquid medium, this reporter became induced during the transition to stationary phase (Figure 1A), which is consistent with previous reports [12, 14, 27]. The GFP reporter level remains high in the stationary phase, which also explains its initially high fluorescence in our plate reader experiments, where the culture is inoculated using stationary-phase cells. The observed induction of curli expression occurred at similar densities in the cultures with different initial inoculum size. In both cases the onset of induction apparently coincided with the reduction of the growth rate, which might occur due to depletion of amino acids in the medium and induction of the stringent response [36, 37], consistent with the proposed role of the stringent response in the regulatory cascade leading to curli gene expression [18, 23]. In agreement with that, curli expression was strongly reduced when *E. coli* cultures were grown in a concentrated TB medium (Figure S1A) or when TB medium was supplemented with serine (Figure S2A). Moreover, curli reporter induction was enhanced by the addition of serine hydroxamate (SHX), which is known to mimic amino acid starvation and induce the stringent response [38] (Figure S2A).

**Figure 1:**
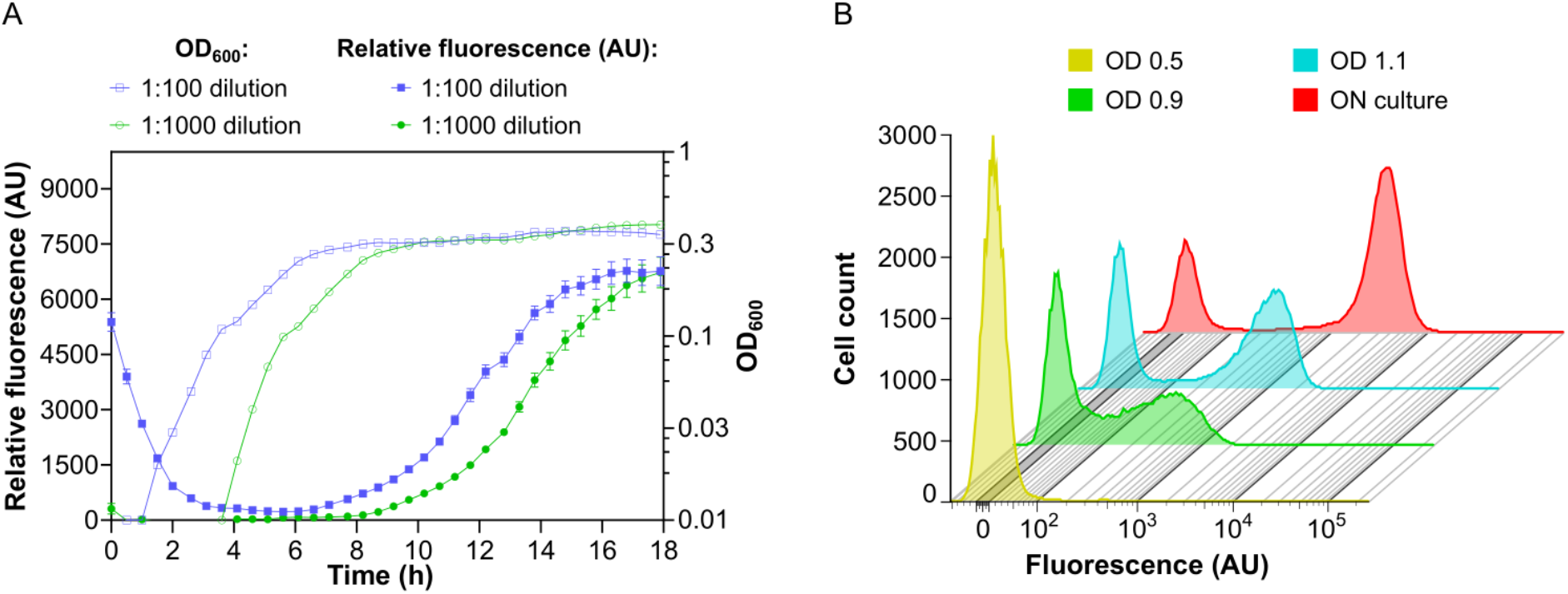
Bimodal activation of curli gene expression in *E. coli* planktonic cultures. *E. coli* cells carrying genomic transcriptional reporter of *csgBAC* operon were grown in liquid tryptone broth (TB) medium at 30 °C under constant shaking. **(A)** Optical density (OD_600_) and relative fluorescence (fluorescence/OD_600_; AU, arbitrary units) of the culture during growth in a plate reader, starting from two different dilutions of the overnight culture. Error bars indicate standard error of the mean (SEM) of 10 technical replicates. **(B)** Distribution of single-cell fluorescence levels, measured by flow cytometry, of cultures grown from overnight culture diluted 1:1000 to indicated OD_600_ or overnight (ON; 25 h) in flasks in an orbital shaker.

In order to investigate whether curli expression was uniform or heterogeneous within planktonic *E. coli* populations, we next measured curli reporter activity in individual cells using flow cytometry. The reporter was induced only in a fraction of cells, and this bimodality of curli expression became increasingly more pronounced at later stages of culture growth, reaching its maximum in the overnight culture (Figure 1B). Thus, the bimodal induction of curli gene expression is observed not only in biofilms but also in well-mixed planktonic cultures. Upon inspection of the flask culture by microscopy, no cell aggregation could be observed, suggesting that curli gene induction in a subpopulation of bacteria is not due to the formation of suspended biofilm-like aggregates [39]. While curli activation was more pronounced in a cell culture growing in an orbital shaker in a flask (Figure 1B), bimodality was also observed for cultures grown in a plate reader (Figure S1B and Figure S2B). Notably, stimulation of curli expression by SHX, or its suppression by additional nutrients, affected the fraction of positive cells rather than their expression levels (Figure S1B and Figure S2B).

### Bimodality of curli gene expression does not require the known c-di-GMP regulatory network

Subsequently, we investigated dependence of bimodal activation of curli gene expression in planktonic culture on the known regulation of curli expression in *E. coli* by the global (DgcE and PdeH) and local (DgcM and PdeR) pairs of DGC/PDE enzymes [12, 14, 22] (Figure 2A). In agreement with this model of c-di-GMP-dependent regulation, activation of curli reporter was completely abolished in the absence of MlrA (Figure 2B), and it was strongly reduced by deletions of *dgcE* and *dgcM* and enhanced by deletions of *pdeH* and *pdeR* genes (Figure 2C). Notably, under these conditions a small fraction of curli-positive cells could still be detected in cultures of both DGC gene deletion strains, and a small fraction of curli-negative cells was observed for both PDE gene deletion strains, indicating that deletions of these genes do not eliminate the bimodality of curli gene expression or its levels in curli-positive cells but rather change the probability of curli genes to become activated in individual cells.

**Figure 2:**
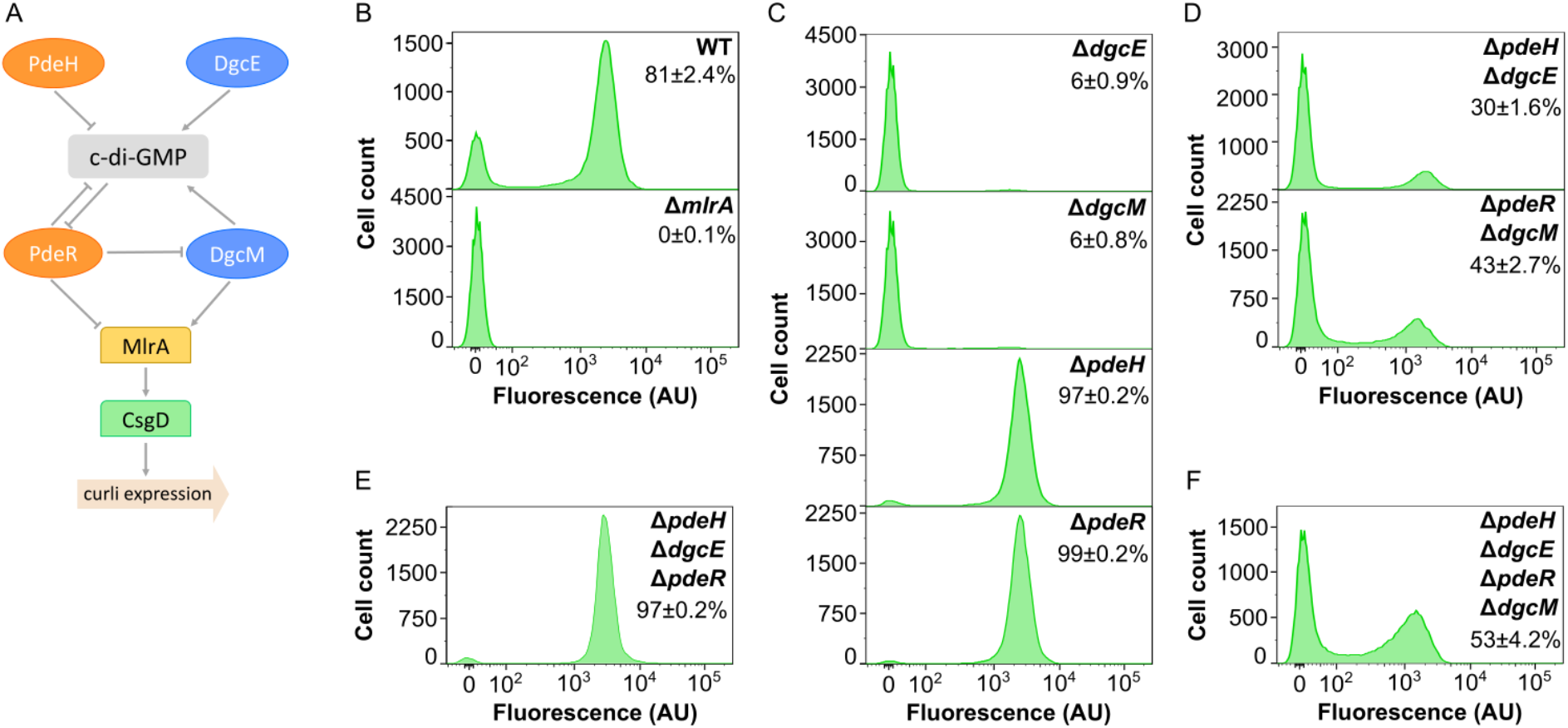
Regulation of curli gene expression by c-di-GMP. **(A)** Current model of regulation of curli gene expression by c-di-GMP in *E. coli*, adapted from [27]. The regulation is mediated by two pairs of diguanylate cyclases (DGCs; blue) and phosphodiesterases (PDEs; orange). PdeH and DgcE control global levels of c-di-GMP, whereas PdeR and DgcM mediate local c-di-GMP-dependent regulation of curli gene expression by controlling activity of transcription factor MlrA, which activates another curli-specific transcription factor CsgD. **(B-F)** Flow cytometry measurements of curli gene expression in *E. coli* planktonic cultures grown in TB overnight in flasks in an orbital shaker, shown for the wild-type (WT) and Δ*mlrA* strain (B), and individual (C), double (D), triple (E) and quadruple (F) deletions of DGC or PDE enzymes, as indicated. Fraction of positive cells in the population (mean of three biological replicates ± SEM) is indicated for each strain. Note that the scale in the *y* axes is different for individual strains to improve readability.

Consistently, bimodality was also retained upon combined deletions of pairs of DGC/PDE genes. Removal of the entire global level of c-di-GMP regulation led to a bimodal pattern of curli reporter activation within the population of Δ*pdeH* Δ*dgcE* cells (Figure 2D) that was similar to that observed in the wild-type culture, despite a smaller fraction of curli-positive cells in the deletion strain. Even more surprisingly, the distribution of curli reporter expression remained bimodal upon removal of the local level of c-di-GMP regulation in Δ*pdeR* Δ*dgcM* strain, although the fraction of curli-positive cells was reduced in this background, too.

We observed that level of curli gene expression in Δ*pdeR* Δ*dgcM* strain was lower than in Δ*pdeR* strain, which confirms that DgcM can promote curli expression independently of PdeR [12, 14, 22] (Figure 2A). This conclusion is further supported by the comparison between the triple deletion strain Δ*pdeH* Δ*dgcE* Δ*pdeR* that expresses only DgcM (Figure 2E) and the quadruple deletion strain Δ*pdeH* Δ*dgcE* Δ*pdeR* Δ*dgcM* that lacks the entire regulatory network (Figure 2F), with much higher fraction of culri-positive cells observed in the former background. Most importantly, the bimodality of curli expression was still observed in the quadruple deletion strain (Figure 2F).

We thus conclude that, whereas the known c-di-GMP-dependent regulation of MlrA activity does clearly affect the fraction of curli-positive cells, it is not required to establish bimodality of curli gene expression in the planktonic cell population. The network has further only little impact of the level of curli reporter activity in individual positive cells (i.e., on position of the positive peak in the flow cytometry data), although reporter intensity in positive cells appeared to be slightly reduced in the strains lacking DgcM.

*Vibrio cholerae* transcriptional regulator VpsT, a close homologue of CsgD, has been shown to be directly regulated by binding to c-di-GMP [40]. Furthermore, in both *S.* Typhimurium and *E. coli*, curli gene expression might also be regulated by c-di-GMP independently of *csgD* transcription [41–43]. We thus aimed to examine whether *E. coli* curli gene expression was no longer sensitive to the global cellular level of c-di-GMP in the absence of the local PdeR/DgcM regulatory module. Indeed, whereas the overexpression of DgcE or of PdeH, which both affect the global pool of c-di-GMP in *E. coli*, had strong impacts on the fraction of curli-positive cells in the wild type, the quadruple mutant was insensitive to such overexpression (Figure 3). This result suggests that in this background, the expression of the *csgBAC* reporter is indeed no longer affected by global c-di-GMP levels. This further excludes regulation by other DGCs that contribute to the common c-di-GMP pool, although it does not rule out hypothetical regulation by a local DGC/PDE pair that is insensitive to global c-di-GMP levels.

**Figure 3:**
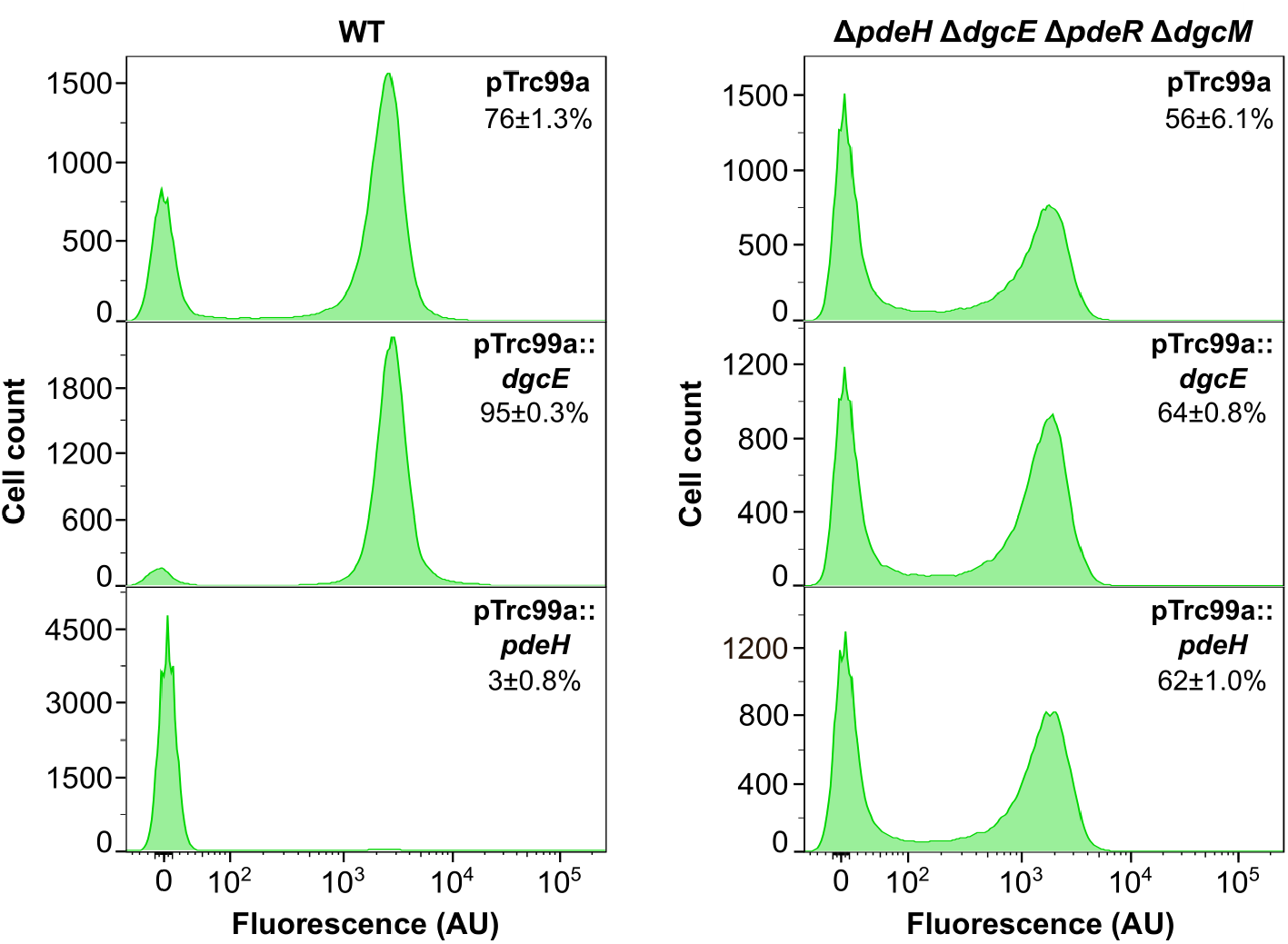
Decoupling of curli gene expression from c-di-GMP regulation in the absence of PdeR/ DgcM regulatory module. *E. coli* wild-type (WT) cells or cells lacking c-di-GMP regulatory enzymes (Δ*pdeH* Δ*dgcE* Δ*pdeR* Δ*dgcM*) were transformed with either empty pTrc99a plasmid (control) or with pTrc99a plasmid carrying *dgcE* (pTrc99a∷dgcE) or *pdeH* (pTrc99a∷pdeH) genes. Expression from the vector was induced with 1 μM IPTG. Bacteria were grown in TB overnight in flasks with shaking and cultures were subjected to the flow cytometry analysis. Fraction of positive cells in the population (mean of three biological replicates ± SEM) is indicated for each strain.

### Curli gene activation shows higher growth-condition dependent variability in the absence of the c-di-GMP regulatory network

We next explored how the fraction of curli-positive cells in the population depends on the conditions of culture growth, with or without regulation by c-di-GMP. As noted above, when the wild-type *E. coli* culture was grown in multi-well plates, both the fraction of curli-positive cells and the level of reporter activity in positive cells were moderately reduced compared to incubation in a flask in an orbital shaker (Figure 2B, Figure 4A and Figure S1B). The reduction in the number of curli-positive cells under these growth conditions was even more pronounced for Δ*pdeR* Δ*dgcM* or Δ*pdeH* Δ*dgcE* Δ*pdeR* Δ*dgcM* strains (Figure 4A and Figure S3), where only a small fraction of cells (12%) became positive, compared to 40-50% in the flask culture (Figure 2D,F). Interestingly, this difference was less for the Δ*pdeH* Δ*dgcE* strain (24% vs 30%). Deletions of individual DGC or PDE genes generally showed expected phenotypes, but fractions of curli-positive cells in Δ*pdeH* and Δ*pdeR* strains were again reduced compared with flask cultures (Figure 2C). Another notable difference between flask and multi-well plate cultures was that the low-fluorescence peak of the wild-type culture was not fully negative but apparently contained a large fraction of cells with incompletely activated curli reporter, which could also be seen in the Δ*pdeH* or Δ*pdeR* strains but not in the Δ*pdeH* Δ*dgcE* Δ*pdeR* Δ*dgcM,* Δ*pdeR* Δ*dgcM* or Δ*pdeH* Δ*dgcE* strains. Similar results were obtained even upon prolonged incubation in the plate reader (Figure S4), confirming that the observed difference with the overnight flask culture was not because of the different growth stage but rather due to differences in growth conditions.

**Figure 4:**
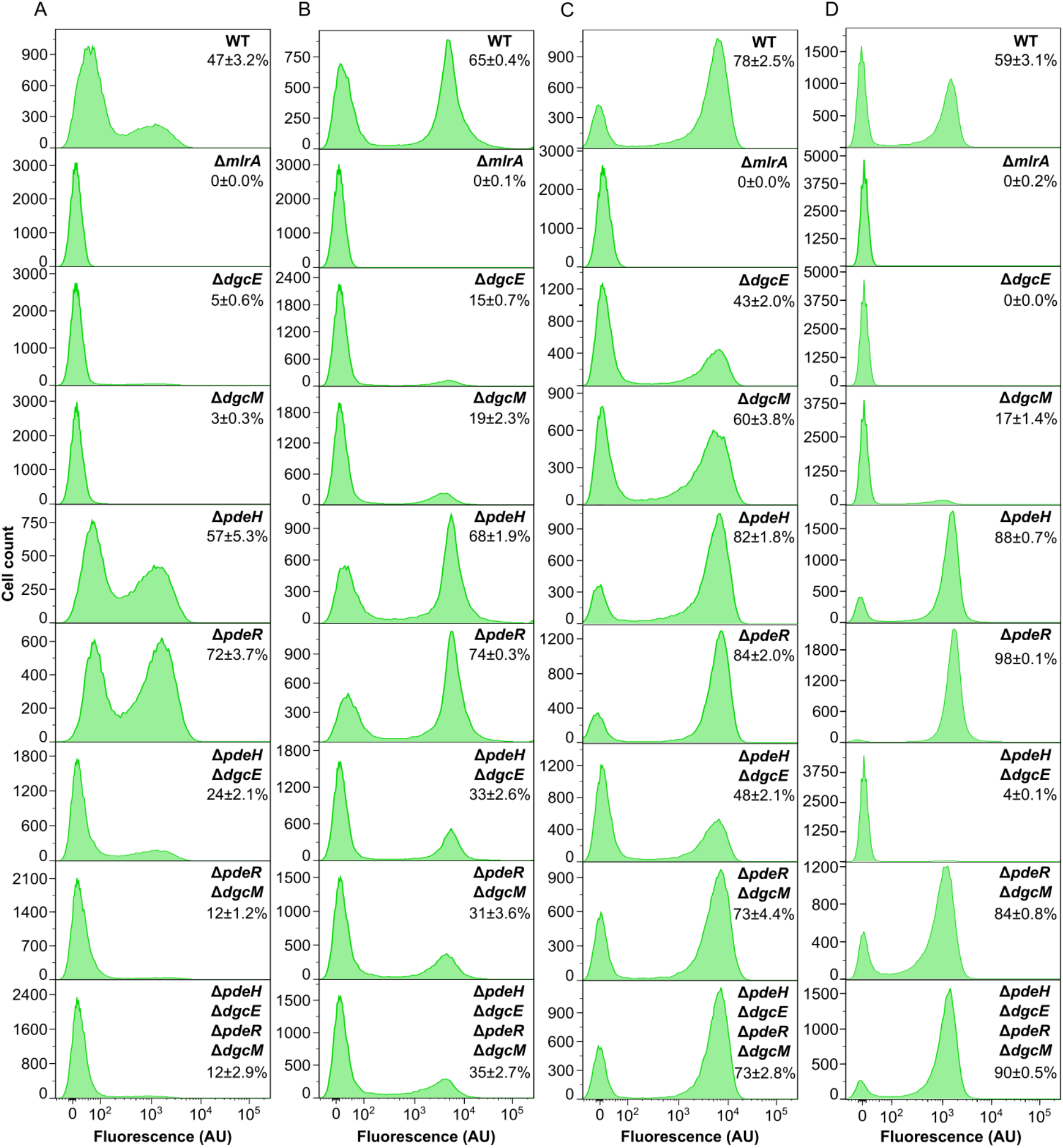
Dependence of curli gene expression on c-di-GMP regulation under different growth conditions. **(A-D)** Flow cytometry measurements of curli expression in the wildtype (WT) and indicated gene deletion strains either after 24 h of growth in liquid TB in a plate reader (A) or as submerged biofilms without agitation (B), as macrocolony biofilms on salt-free LB agar (C), or as overnight planktonic cultures in salt-free LB in flasks in an orbital shaker. Fraction of positive cells in the population (mean of three biological replicates ± SEM) is indicated for each strain.

We further tested reporter activation under growth conditions that favor biofilm formation. During formation of static submerged biofilms in multi-well plates, where cultures are grown without shaking, the overall curli activation pattern in cell populations of individual or double and quadruple deletion strains (Figure 4B) was comparable to that in the flask culture grown in the orbital shaker (Figure 2B-D, F), although effects of DGC and PDE gene deletions on the fraction of curli-positive cells were apparently reduced. Curli gene activation in individual mutant strains correlated well with the levels of submerged biofilm formation (Figure S5), but the lack of regulation by c-di-GMP resulted in stronger reduction of the biofilm biomass than simply expected from its effect on curli gene expression, consistent with additional roles of c-di-GMP in biofilm formation.

All strains were also grown in the form of macrocolony biofilms on agar plates containing the salt-free LB medium, as done previously [26] (Figure 4C), as well as in the liquid salt-free LB medium as a control (Figure 4D). Expression of curli reporter in individual cells within the macrocolony was analyzed by flow cytometry immediately after colony resuspension (see Materials and Methods), so that the level of stable GFP reporter should reflect cell-specific curli expression within the macrocolony. The overall dependency of reporter activation on the deletions of individual DGC and PDE genes was again qualitatively similar to other growth conditions (Figure 4C). However, the fraction of curli-positive cells in the macrocolony biofilms was less sensitive to deletions of individual *dgc* or *pdh* genes than in planktonic cultures, regardless whether the latter were grown in TB (Figure 2C) or in the salt-free LB (Figure 4D). This agrees with stronger Congo red staining of macrocolonies of Δ*dgcE* and Δ*dgcM* strains compared to the Δ*mlrA* negative control (Figure S6). Thus, during macrocolony biofilm formation, regulation by c-di-GMP appears to be less important for the activation of curli gene expression, possibly because other regulators contribute to this activation in a highly structured environment. Importantly, the curli-positive cell fraction in both macrocolony and planktonic cultures of Δ*pdeR* Δ*dgcM* and Δ*pdeH* Δ*dgcE* Δ*pdeR* Δ*dgcM* strains growing on the salt-free LB was much higher than under other growth conditions.

To summarize, under all of the tested growth conditions, regulation by the known c-di-GMP regulatory network was not required for bimodal curli activation, although it indeed affected the fraction of curli-positive cells. Nevertheless, the absence of this c-di-GMP control apparently makes the fraction of curli-positive cells in the population more sensitive to environmental conditions, since this fraction exhibited much larger variation in the quadruple deletion compared to the wild-type strain among different growth conditions tested here.

### Dynamics and variability of stochastic curli gene activation in individual cells is controlled by the c-di-GMP regulatory network

In order to investigate the dynamics of curli gene activation, and the impact of c-di-GMP regulation, at the single-cell level, we utilized a “mother machine” device — a microfluidic chip where growth of individual bacterial cell lineages could be followed in a highly parallelized manner over multiple generations [35] (Figure 5 and Supporting protocols). Since our design of the mother machine allows rapid switching of the medium supplied to the cells (Figure 5D), we could mimic nutrient depletion in order to activate curli expression in continuously growing single cells. For that, wild-type, individual DGC or PDE, or quadruple deletion strains were first loaded into the mother machine from overnight cultures, allowed to grow in fresh TB medium for several generations, and then shifted to the TB medium that was pre-conditioned by growing a batch culture (see Materials and Methods and Supporting protocols). A fraction of curli-positive cells was observed at the beginning of these experiments, which was strain-specific and consistent with the expected fraction of positive cells in the overnight cultures of each respective strain. After resuming exponential growth in the fresh medium all cells turned off curli expression (Figure 6A,C, Figure S7 and Movie S1, Movie S2 and Movie S3). Following the shift to conditioned medium, cell growth rate was strongly reduced (Figure 6E and Figure S8), and after several generations of slow growth, individual cells of all strains activated curli expression, while other cells remained in the curli-off state (Figure 6A,C,F, Figure S7 and Movie S1, Movie S2 and Movie S3). Importantly, the frequencies of curli-positive cells in individual mutants were consistent with those of the planktonic cultures, confirming that the overall control of curli gene expression by the c-di-GMP regulatory network is similar for growth in conditioned medium in the mother machine.

**Figure 5:**
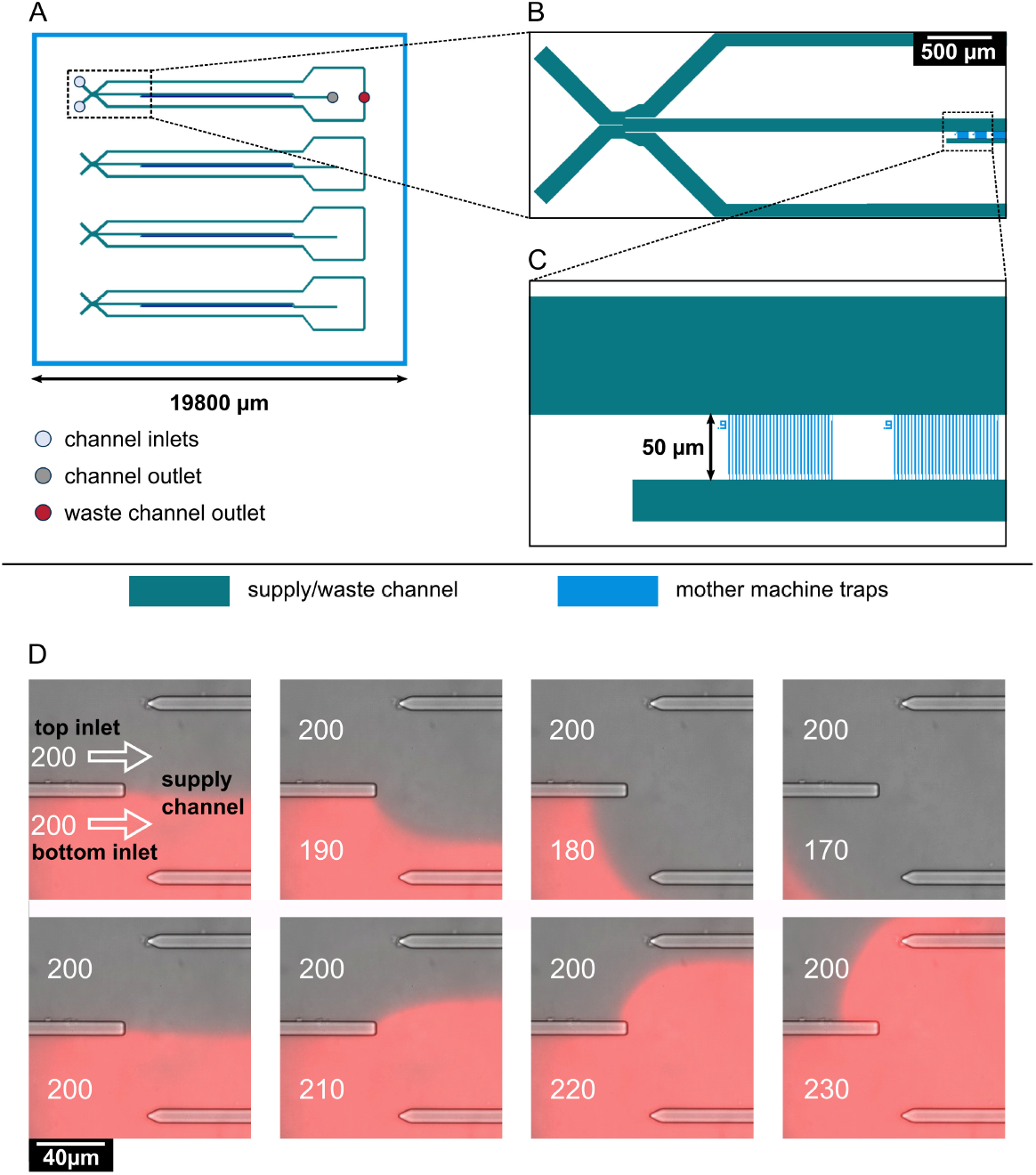
Design and operation of the microfluidic mother machine device. **(A)** Schematic overview of the chip layout, featuring four independent supply channels (green) for cell inoculation and media supply. **(B)** Detailed view of the area marked by a rectangle in (A), showing the switching junction and a part of the mother machine cultivation sites (blue). The junction is formed by two inlets, leading to one central supply channel. Control of pressure at each inlet allows for prioritization of one medium over the other through the supply channel, and, ultimately, the mother machine cultivation sites. Residual medium flows out through waste channels located to each side of the central supply channel. Medium flowing through the supply channel exits the chip through one outlet. **(C)** Detailed view of the area marked in **(B)** by a rectangle, showing the mother machine cultivation sites. Each of the four channels contains 57 mother machine cultivation sites, each of which contains 30 mother machine traps with widths of either 0.9, 1, or 1.1 μm. The mother machine traps feature a 0.3 μm wide constriction on the bottom, preventing the mother cell from exiting the trap while allowing perfusion of the medium. The supply channels (green) are 8 μm in depth, the mother machine traps (blue) are 0.8 μm in depth. **(D)** On-chip medium switching visualized by merged phase contrast and mCherry images of the channel junction. Media are supplied through separate inlets (top and bottom), which are separated in the center of the channel by a PDMS barrier. The direction of flow is indicated by white arrows. Water was supplied through the top inlet, while a 0.2 μM sulforhodamine B solution was supplied through the bottom inlet, visualizing the flow pattern in the junction. The pressure at the top inlet was kept constant at 200 mbar. Depending on the pressure set at the bottom inlet, it is possible to select which one of the two media flows into the central supply channel to the mother machine growth sites.

**Figure 6:**
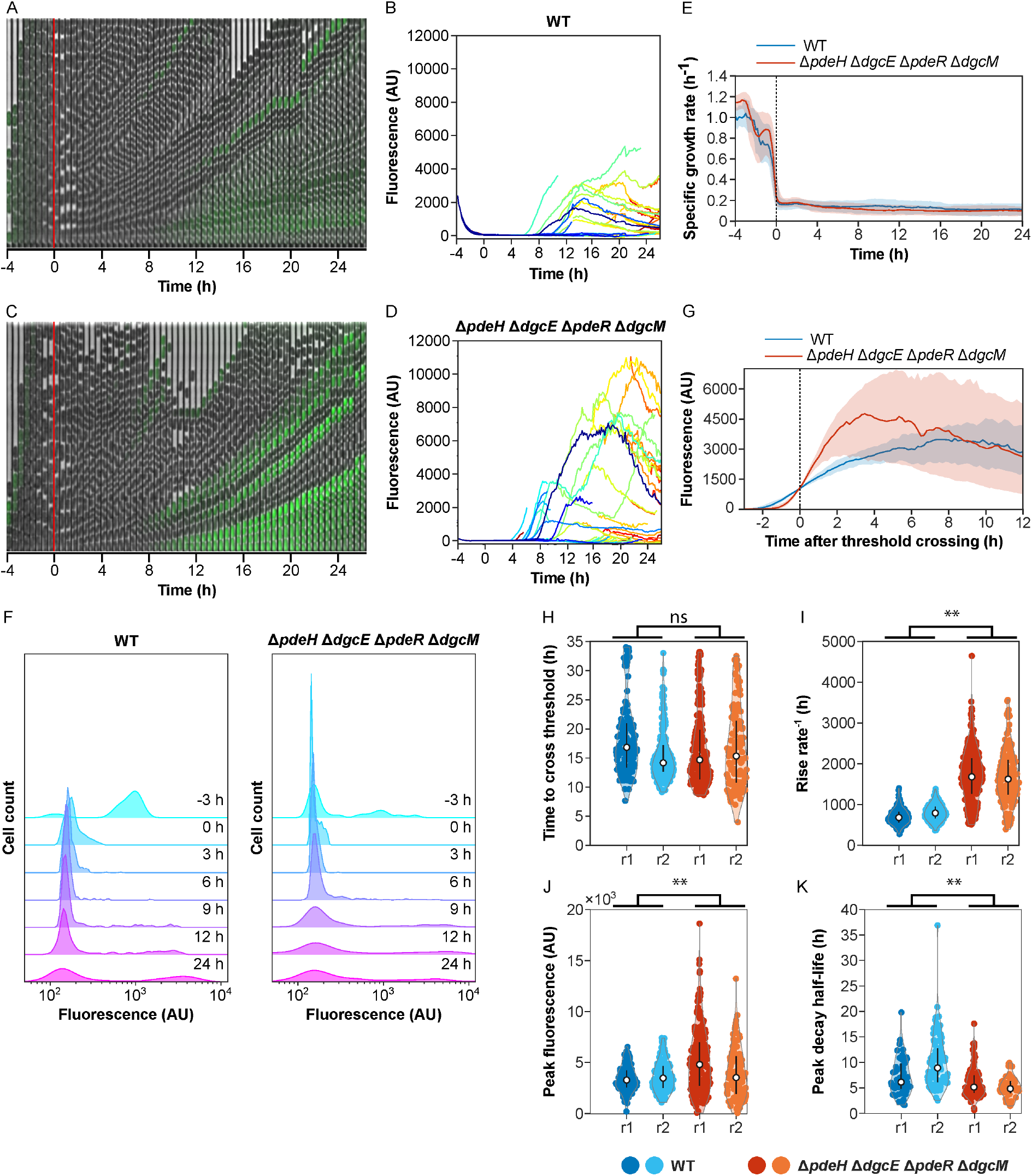
Impact of c-di-GMP regulation on dynamics of curli gene induction in individual cells. *E. coli* cells in a microfluidic device (mother machine) were shifted from a fresh to conditioned TB medium after 4 h of growth to induce curli expression. **(A-D)** Examples of image time series and single-cell fluorescence traces for the wild-type (WT) (A,B) and for the Δ*pdeH* Δ*dgcE* Δ*pdeR* Δ*dgcM* strain disabled in the c-di-GMP regulation (C,D), growing in a mother machine in one experiment. Expression of the curli reporter is indicated by the green overlay on the phase contrast image. **(E)** Median instantaneous growth rate — fold rate of change in length — of cells grown in microfluidics experiments as described in (A). The switch to conditioned medium is at time zero, as indicated. Shaded area is the interquartile range. The number of cells in the device varies with time, but is on average *n* = 296 for WT and *n* = 522 for the quadruple deletion strain. **(F)** Distributions of curli expression at different time points in the microfluidics experiment. Shown are kernel density estimates of curli expression in WT and in the quadruple deletion strain at selected time points. **(G)** Median curli expression profile for single-cell traces aligned by the time at which they exceed a threshold of 10^3^ fluorescence units. Shaded area is interquartile range; *n* = 128 for WT and 230 for the quadruple deletion strain. **(H-K)** Distributions of curli induction parameters for the wild-type and for the c-di-GMP-regulation disabled strain, in two independent experiments, r1 (same experiment as in A-G) and r2. Shown are histograms of the times at which a threshold of 10^*3*^ fluorescence units were crossed **(H)**, the maximum rates of increase in fluorescence **(I)**, the fluorescence amplitudes at the first peak **(J)** and the half-time of fluorescence decay after reaching the peak **(K)** for cell traces from both microfluidics experiments. \*\**p*<10^−2^ in an unpaired two-sample *t*-test assuming normal distributions with unknown and unequal variances. ns, non-significant.

Although the overall frequency of curli reporter activation was comparable for the wild-type and Δ*pdeH* Δ*dgcE* Δ*pdeR* Δ*dgcM* cells lacking c-di-GMP regulatory network, we further compared the dynamics of single-cell curli activation in both backgrounds. First, we observed that, in both strains, curli reporter activation was transient — and after several generations in the curli-on state individual cells turned curli expression off again during continuous growth in the conditioned medium (Figure 6B,D, Figure S8 and Figure S9). In some cases, the initial transient pulse of activation was even followed by a second activation event. Although such transient curli reporter activation was observed in both backgrounds, with a similar delay after exposure to conditioned medium (Figure 6H), its dynamics showed several important differences between strains. Most clearly, the rate of curli activation in individual cells was higher in the absence of the c-di-GMP-regulatory network (Figure 6G,I and Figure S10A,B), which led to stronger but more transient increase in their fluorescence (Figure 6J,K and Figure S10B). Additionally, the level of reporter induction showed greater intercellular heterogeneity in the Δ*pdeH* Δ*dgcE* Δ*pdeR* Δ*dgcM* strain (Figure 6,D,J, Figure S9). Thus, the control of curli expression by c-di-GMP reduces the rate but also the cell-to-cell variability of curli induction within population.

## Discussion

Expression of the curli biofilm matrix genes is known to be heterogeneous or even bistable in communities of *E. coli* [25–28] and *S.* Typhimurium [29, 30], which might have important functional consequences for the biomechanics of bacterial biofilms [27] and for stress resistance and virulence of bacterial populations [29, 30]. In the well-structured macrocolony biofilms, differentiation into subpopulations with different levels of curli matrix production is largely deterministic and driven by gradients of nutrients and oxygen [18], but bimodality of curli expression might also emerge stochastically, in a well-mixed population or between cells within the same layer of a macrocolony [27, 29, 30]. How this bimodality originates within the extremely complex regulatory network of curli genes [17, 19, 23, 27] remains a matter of debate. Although earlier work proposed that bistable expression of curli might be a consequence of positive transcriptional feedback in *csgD* regulation [30, 44], the most recently suggested model attributed bistability to the properties of the c-di-GMP regulatory switch formed by DgcM, PdeR and MlrA [27, 31]. Furthermore, although this bimodal pattern of curli gene expression has typically been referred to as bistable, temporal dynamics and stability of its induction in individual cells has not been directly investigated.

Here, we studied the expression of the major curli *csgBAC* operon in well-stirred planktonic *E. coli* cultures as well as in submerged and macrocolony biofilms and during growth in a microfluidics device. Consistent with previous studies [12, 14, 27], the induction of curli gene expression in growing *E. coli* cell populations was observed during entry into the stationary phase of growth. Curli activation was apparently dependent on depletion of nutrients, most likely amino acids, since it could be suppressed by increasing the levels of nutrients or, more specifically, by addition of serine, and conversely, it could be enhanced by the SHX-mediated stimulation of the stringent response. We further observed that activation of the *csgBAC* operon exhibited strongly pronounced bimodality under all tested conditions, resulting in two distinct, curli-positive and -negative, subpopulations of cells even in a well-stirred planktonic culture. Stochastic bimodality of curli reporter activation was confirmed in a microfluidic device, where, following shift to conditioned medium, only a fraction of the cell population turned on curli gene expression.

Such differentiation is apparently consistent with previous reports of bimodal *csgD* expression in the stationary phase of *S.* Typhimurium [29, 30] and *E. coli* [27] culture growth. However, in contrast with the previous interpretation of such a *csgD* expression pattern as bistability, we observed that activation of curli gene expression during continuous cell growth in the microfluidic device was only transient and unstable. Moreover, our data indicate that the bimodality of the *csgBAC* expression is unlikely to be (fully) explained by the bimodal expression of *csgD*. Firstly, compared to the bimodality of the *csgBAC* reporter, the previously reported bimodality of *csgD* expression in *E. coli* [27] was much less pronounced and became apparent only in later stages of planktonic culture growth. Moreover, in that study, all wild-type cells showed activation of the *csgD* reporter, differing only in the level of this activation, which itself cannot explain the on/off bimodality of the *csgBAC* reporter activation.

Secondly, and most importantly, whereas the bimodality of *csgD* expression was observed to be dependent on regulation by the DgcE/PdeH/DgcM/PdeR network [27], this was clearly not the case for *csgBAC* reporter expression. Under all tested conditions, including planktonic cultures, submerged and macrocolony biofilms, and growth in the microfluidic device, the differentiation into distinct subpopulations of positive and negative cells in our experiments occurred in the absence of this entire c-di-GMP regulatory network. Since removal of the DgcM/PdeR module further made curli gene expression insensitive to global levels of c-di-GMP, contributions of other *E. coli* DGCs and PDEs to curli regulation in this deletion background are unlikely, even if these enzymes might in principle affect curli gene expression in wild-type *E. coli* cells via the common c-di-GMP pool. We therefore conclude that c-di-GMP regulation might be dispensable for activation or bimodality of curli gene expression in *E. coli*, although we could not rule out regulation of curli gene expression by a hypothetical local DGC/PDE module that is insensitive to global c-di-GMP levels.

Nevertheless, c-di-GMP control clearly plays an important role during the establishment of bimodality. Consistent with previous studies, we observed that the DGC and PDE proteins determine the fraction of curli-positive cells [22, 23, 27], although they have no or only little effect on the average level of *csgBAC* expression in individual cells. Our results further suggest that the DgcE/PdeH/DgcM/PdeR regulatory network might make curli expression in *E. coli* populations more robust, helping to ensure that the fraction of curli-positive cells is less variable under different culture growth conditions. This might be related to the observed effect of this network on the temporal expression dynamics in a continuously growing culture, with faster but more heterogeneous activation of curli expression in the absence of the c-di-GMP control, although whether and how there two different kinds of variability and their suppression by are connected remains to be investigated. It is further possible that the increased robustness provided by c-di-GMP regulation is due to the network-induced bimodality of *csgD* expression discussed above [27], which might provide an additional stabilizing feedback.

How might the observed pulsatile activation of the curli-positive state originate at the single-cell level and lead to the differentiation into distinct subpopulations in *E. coli* culture? Pulsing in expression was proposed to be common to many gene regulatory circuits [45, 46, 47]. It was recently described in *E. coli* for the upstream regulator of curli, σ^S^ [48], as well as for the flagellar regulon [49, 50] that is anti-regulated with curli [28]. However, in neither of these cases did pulsing lead to bimodality of expression, and their relation to the observed pulses in curli expression thus remains to be seen.

Regardless of the origin of pulsatile expression, the timing and duration of curli activation pulses can explain how such transient activation leads to differentiation into two distinct subpopulations in the planktonic culture. Since the curli expression is only turned on during transition to the stationary phase, individual cells stochastically activate curli genes just before the culture growth ceases, thus making subsequent reversion to the curli-negative state impossible. Although curli gene regulation in biofilms might be more complex, spatial stratification of gene expression in the macrocolony biofilms was proposed to resemble the temporal regulation of expression in planktonic cultures [18, 26]. Thus, also in this case, curli activation would happen in a cell layer just before it ceases to grow, fixing the differentiation between curli-positive and curli-negative cells within this colony layer.

## Supporting information

Movie S1: Microfluidics experiment showing curli expression in the wild-type E. coli and diguanylate cyclase deletion strains.

Movie S2: Microfluidics experiment showing curli expression in the wild-type E. coli and phosphodiesterase deletion strains.

Movie S3: Microfluidics experiment showing curli expression in the wild-type E. coli and quadruple deletion strain lacking c-di-GMP-regulation.

## Acknowledgements

We thank Sarah Hoch, Silvia González Sierra and Gabriele Malengo for technical help and advice.

## Supporting information

**Figure S1:**
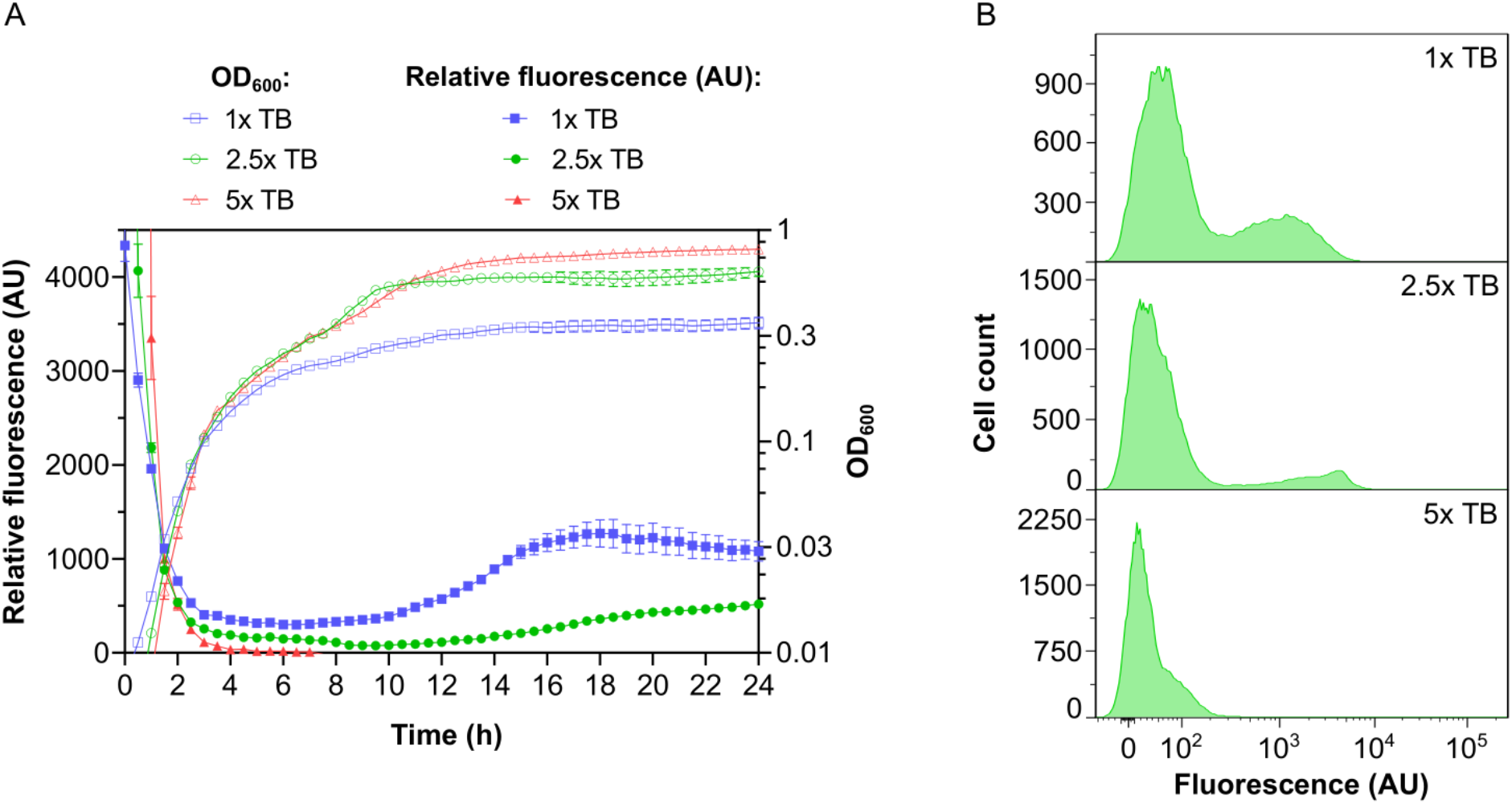
Dependence of curli gene expression on nutrient levels. Wild-type *E. coli* cultures were grown as in Figure 1A but with different indicated concentrations of TB. **(A)** Bacterial growth and activity of transcriptional curli reporter. Error bars indicate SEM of 6 technical replicates. **(B)** Distribution of single-cell fluorescence levels after 24 h of growth in a plate reader measured by flow cytometry. Note that the scale in the *y* axes is different for individual conditions to improve readability.

**Figure S2:**
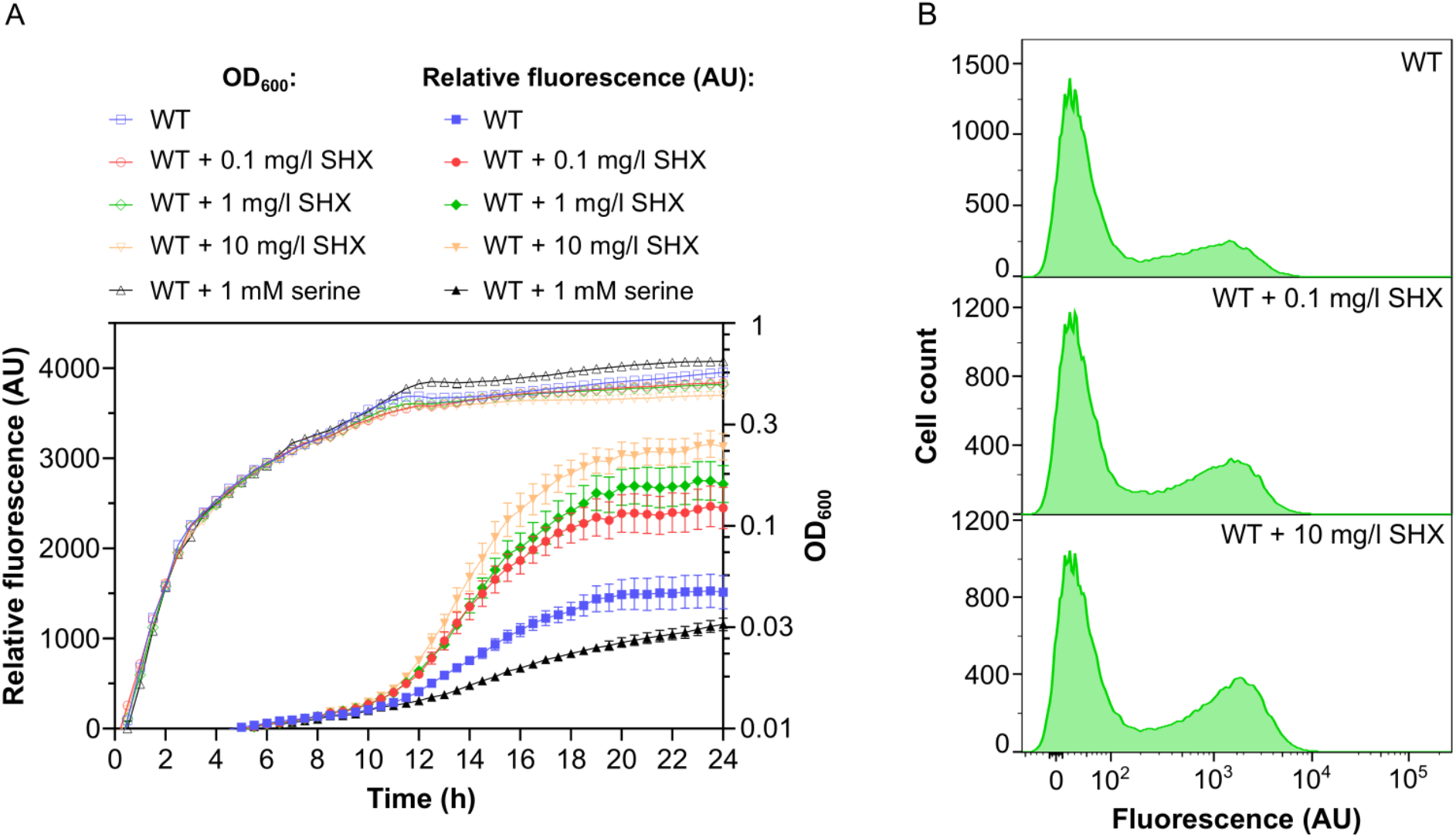
Stimulation of curli gene expression by stringent response. Wild-type *E. coli* cultures were grown as in Figure 1A but with addition of either indicated concentrations of serine hydroxamate (SHX) at inoculation point or 1 mM serine after 6 h of growth. **(A)** Bacterial growth and activity of transcriptional curli reporter. Error bars indicate SEM of 6 technical replicates. **(B)** Distribution of single-cell fluorescence levels after 24 h of growth in a plate reader measured by flow cytometry.

**Figure S3:**
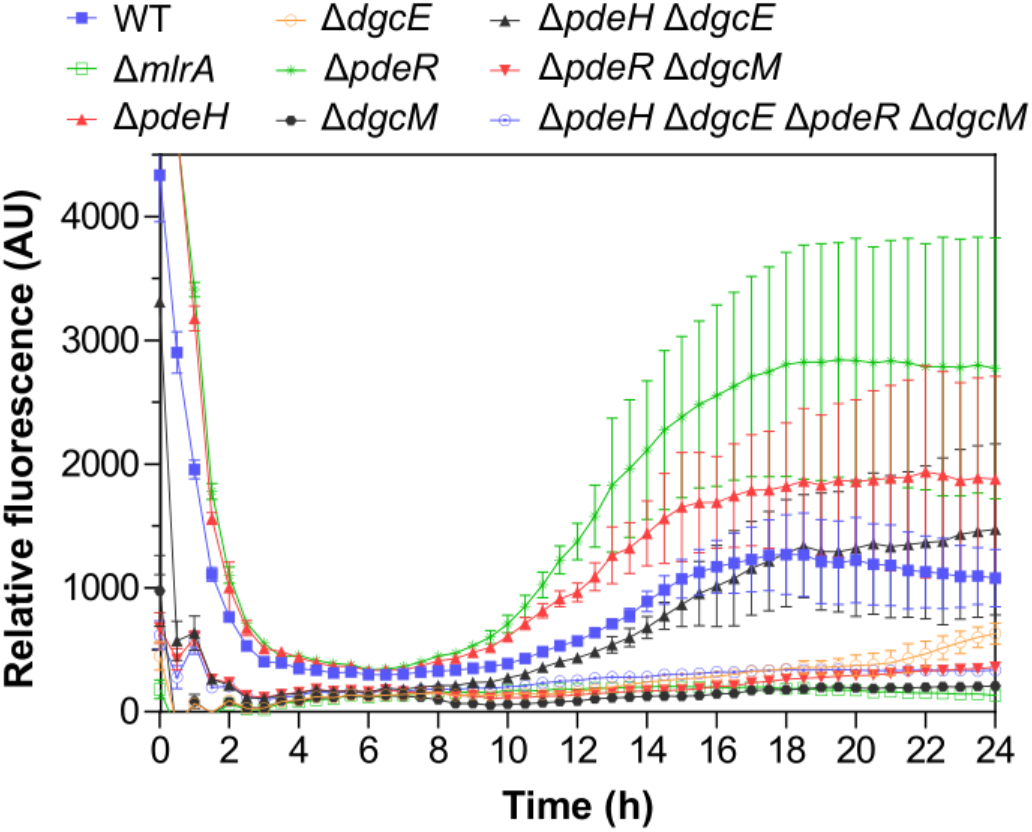
Relative fluorescence of curli reporter in planktonic culture grown in TB medium in a plate reader. Error bars indicate SEM of 5 technical replicates. *E. coli* planktonic cultures of the wild-type (WT), Δ*mlrA* strain, and individual, double and quadruple deletions of DGC or PDE enzymes were grown in TB in a plate reader.

**Figure S4:**
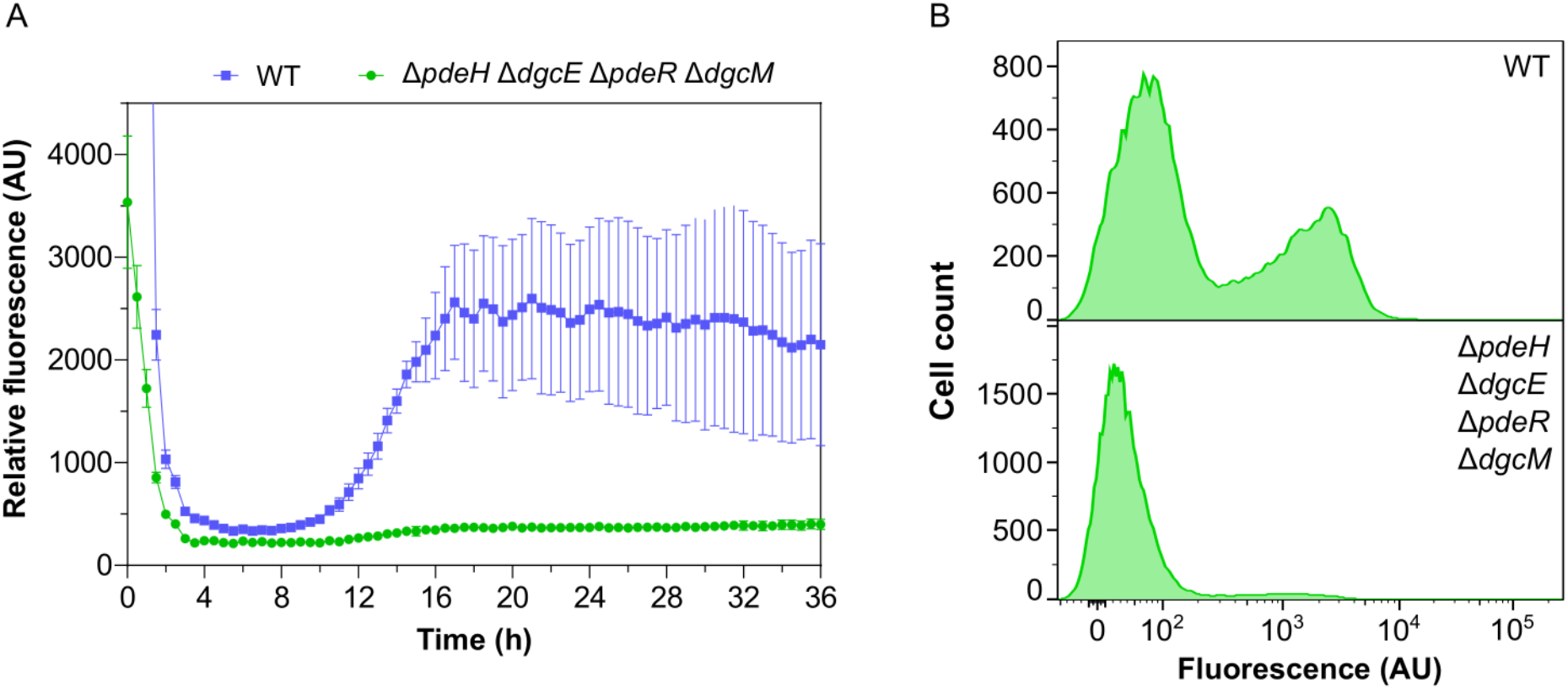
Curli gene expression upon prolonged cultivation in a plate reader. Wild-type (WT) and Δ*pdeH* Δ*dgcE* Δ*pdeR* Δ*dgcM* strains were grown in TB in a plate reader as in Figure S3, but during 36 h. **(A)** Induction of transcriptional curli reporter. Error bars indicate SEM of 10 technical replicates. **(B)** Distribution of single-cell fluorescence levels in populations of both strains after 36 h of growth measured by flow cytometry.

**Figure S5:**
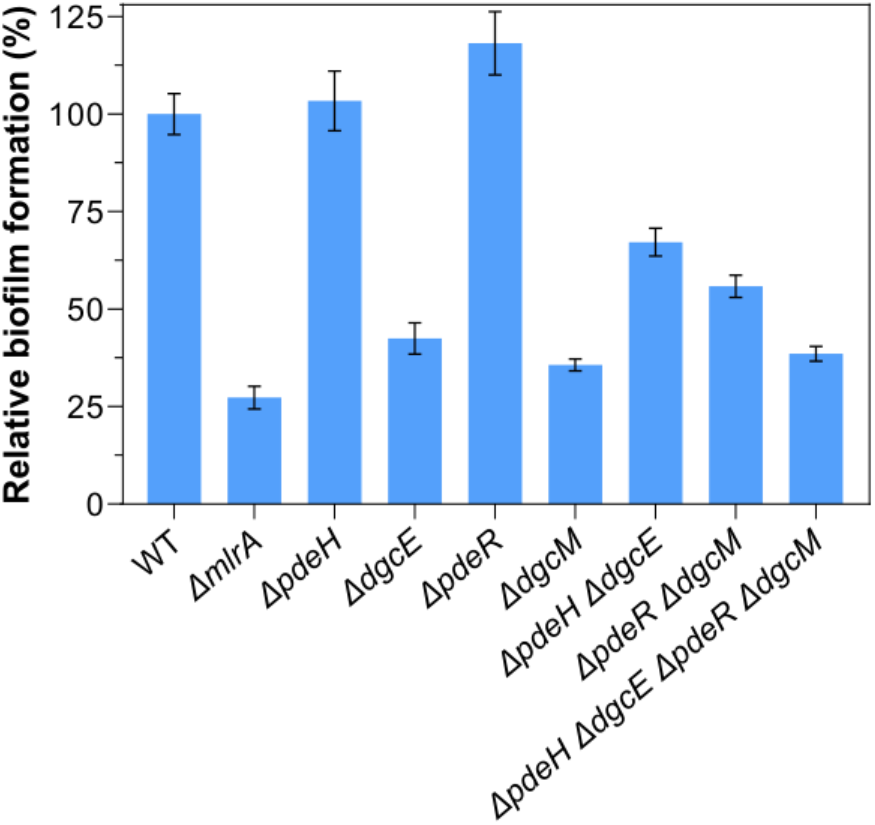
Curli gene expression in submerged biofilm cultures. Biofilm formation by indicated strains grown in multi-well plates in TB medium, quantified using crystal violet (CV) staining. Error bars indicate SEM of 3 independent replicates.

**Figure S6:**
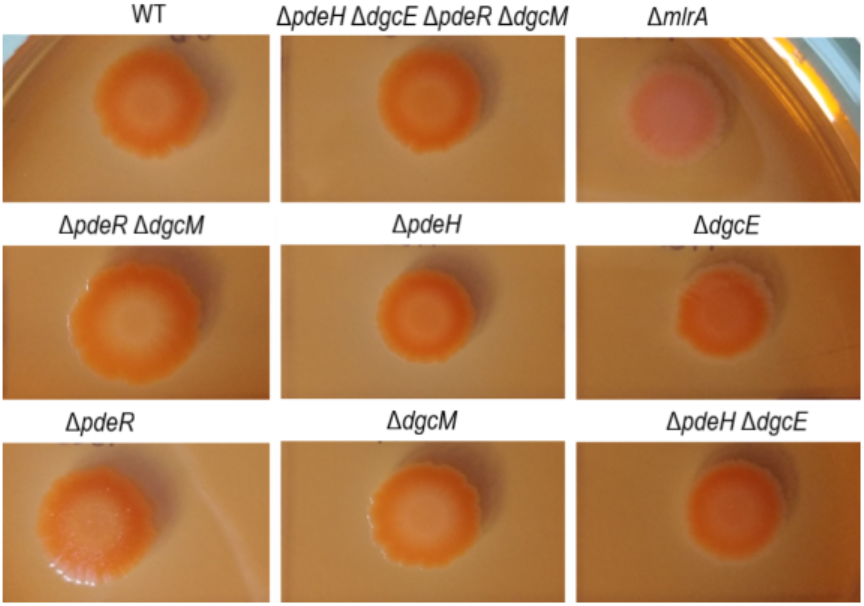
Curli expression in macrocolony biofilms. **I**mages of macrocolonies of indicated strains after 8 days of growth on salt-free LB agar plates.

**Figure S7:**
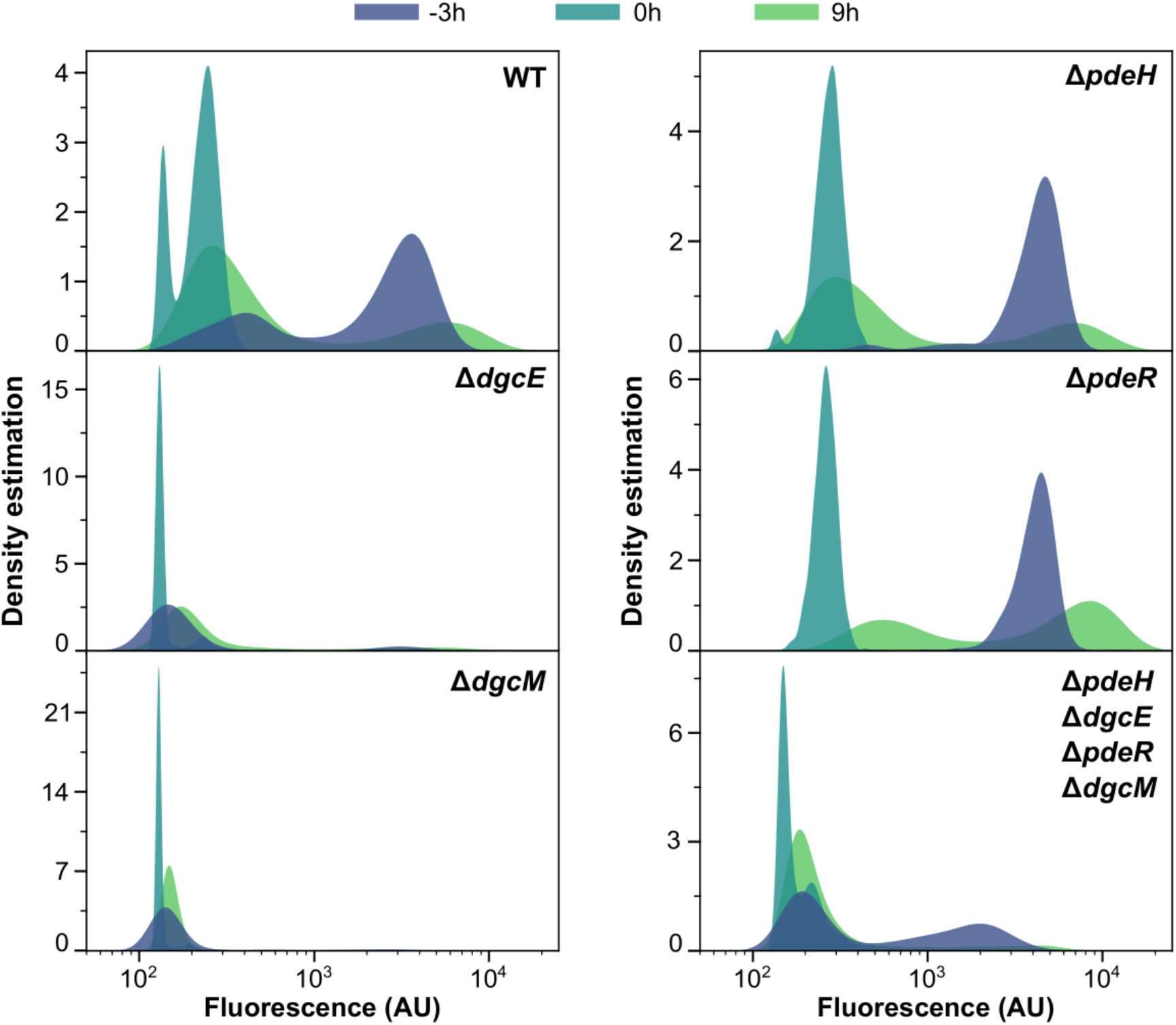
Distributions of curli expression at different time points in the microfluidics experiment. Shown are kernel density estimates of curli expression in the wild type WT, individual and quadruple deletions of DGC or PDE enzymes at selected time points. Cells in stationary phase were loaded into mother machine chip supplied with fresh medium, and then switched to the conditioned media at the ‘’0 hours’’ time point. Note that the scale in the *y* axes is different for individual conditions to improve readability.

**Figure S8:**
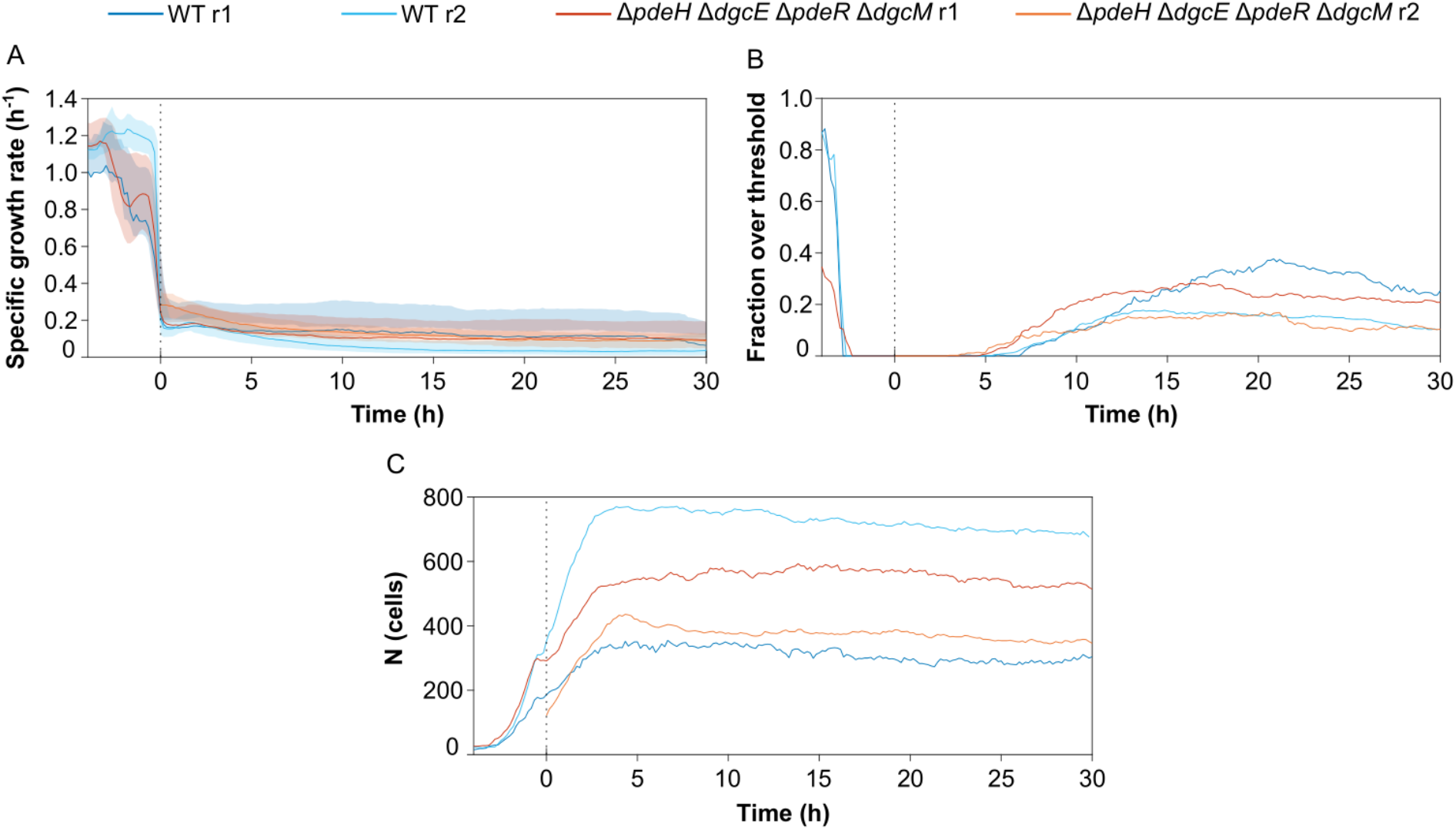
Growth rates and fraction of curli expressing cells over time. Stationary phase cells were introduced into mother machine devices, supplied with fresh medium and then switched to conditioned medium after 4 h of growth, as in Figure 6. **(A)** Median instantaneous growth rates for the wild-type and for the Δ*pdeH* Δ*dgcE* Δ*pdeR* Δ*dgcM* strain disabled in c-di-GMP regulation. Growth rate drops rapidly and cells switch on curli expression after a switch to conditioned medium. Shaded area is interquartile range. **(B)** Fraction of cells with fluorescence exceeding 1000 units. **(C)** Number of detected cells. Two biological replicates (r1 and r2) were performed for each strain; data for the r1 replicate are also shown in Figure 6. Note that in the r2 experiment for the quadruple deletion strain, cells were only imaged after medium switching.

**Figure S9:**
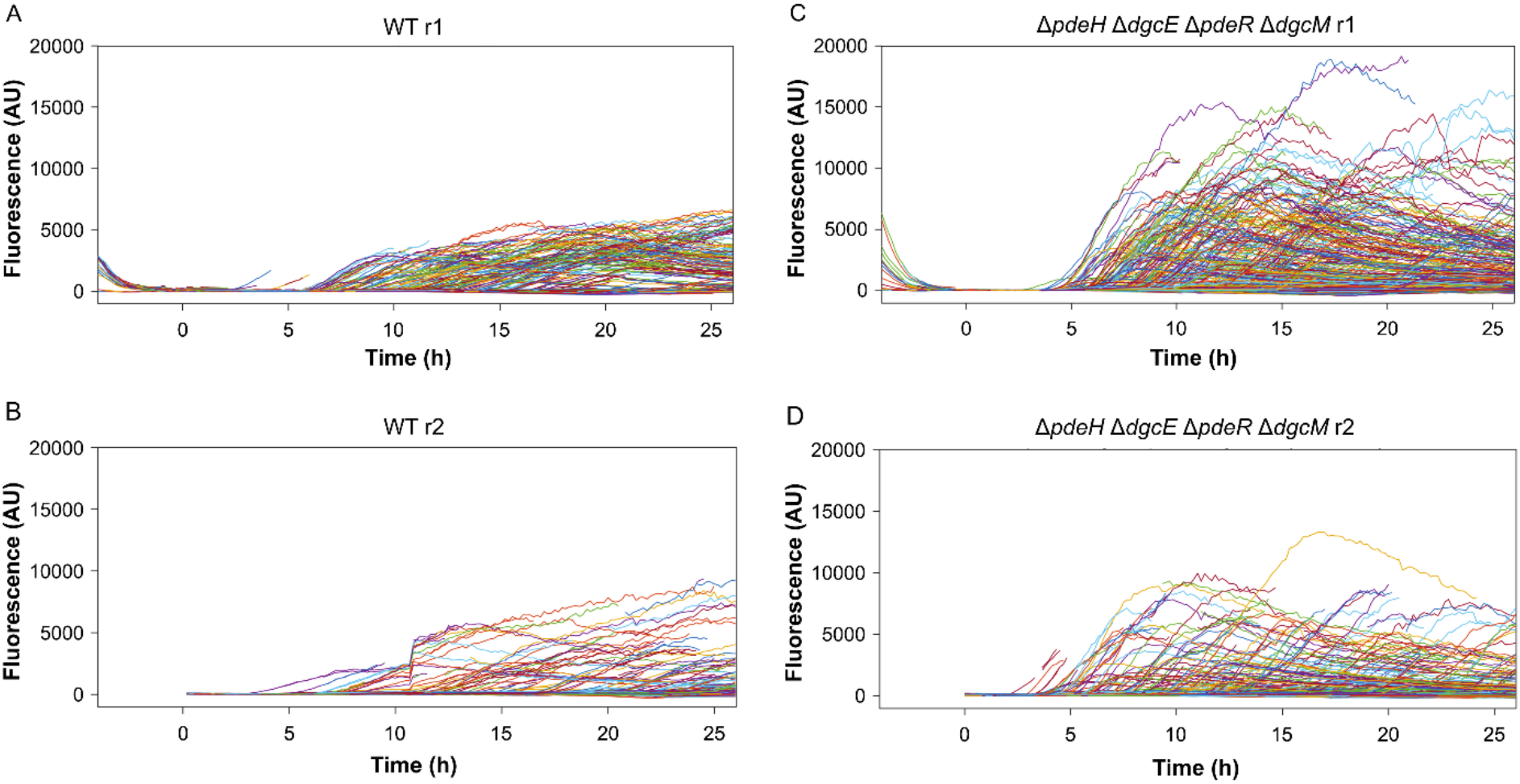
Single-cell traces of cell fluorescence for all cells for the wild-type and for the c-di-GMP-regulation disabled strain. Data are from the same biological replicates (r1 and r2) as in Figure S8.

**Figure S10:**
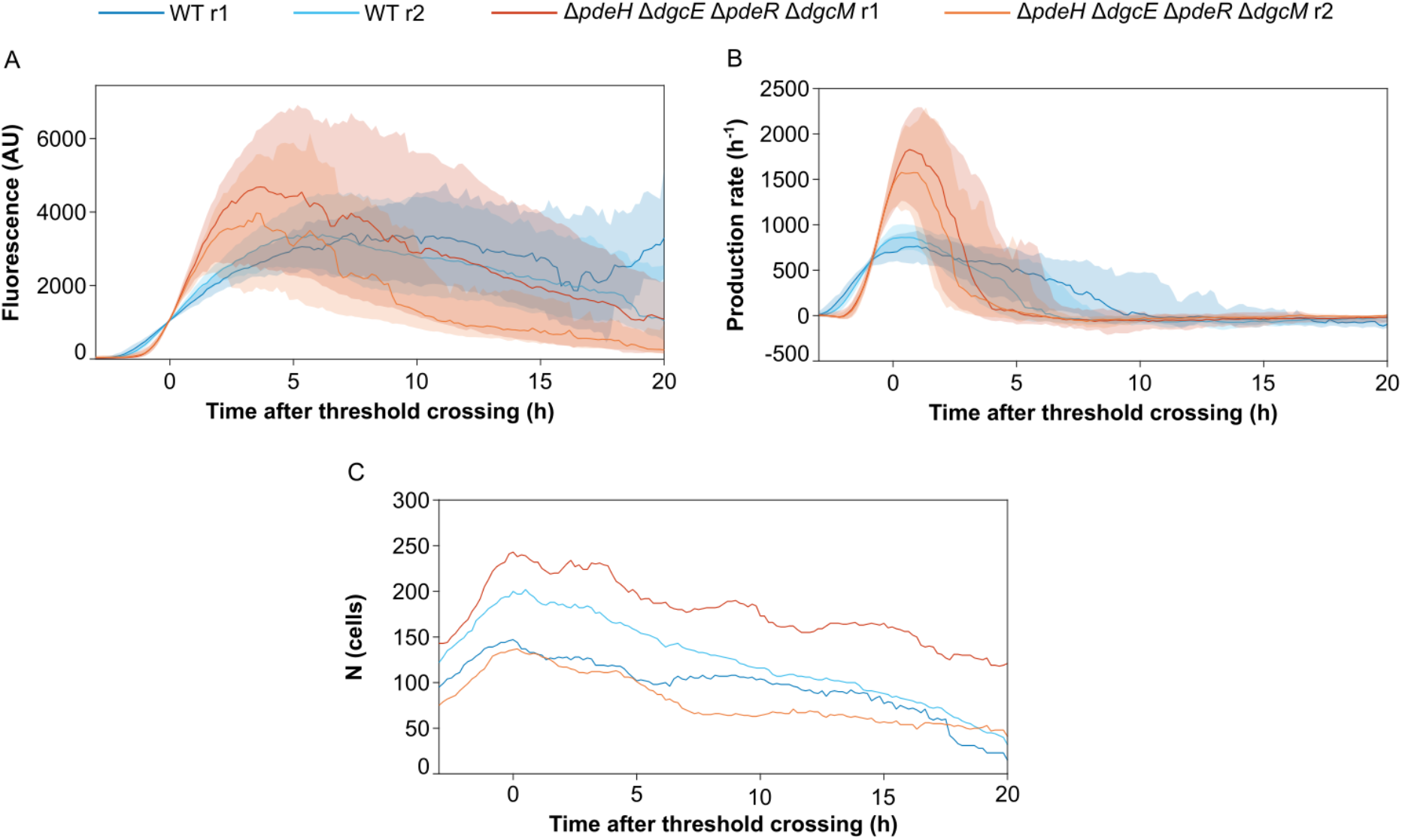
The rate and variability of curli induction for the wild-type and for the c-di-GMP-regulation disabled strain. **(A)** Median curli expression, **(B)** production rate, and **(C)** number of cells for traces from both microfluidics experiments (r1 and r2) aligned by the time at which they exceeded a threshold of 10^*3*^ fluorescence units. Shaded area is interquartile range. Compared to the WT, the rate of curli induction is faster but traces show more variability in a mutant without the global or local c-di-GMP regulatory modules.

**Table S1.** *E. coli* strains and plasmids used in this study.

| Strains | Relevant genotype | Reference |
| --- | --- | --- |
| W3110 | W3110 derivative with functional RpoS | [1] |
| VS1146 | W3110 *csgA::csgA_RBS_sfgfp* | [2] |
| VS1857 | VS1146 $\Delta mlrA$ | This work |
| VS1732 | VS1146 $\Delta pdeH$ | This work |
| VS1720 | VS1146 $\Delta dgcE$ | This work |
| VS1258 | VS1146 $\Delta pdeR$ | This work |
| VS1257 | VS1146 $\Delta dgcM$ | This work |
| VS1717 | VS1146 $\Delta pdeH \Delta dgcE$ | This work |
| VS1713 | VS1146 $\Delta pdeR \Delta dgcM$ | This work |
| VS1885 | VS1146 $\Delta pdeH \Delta dgcE \Delta pdeR$ | This work |
| VS1729 | VS1146 $\Delta pdeH \Delta dgcE \Delta pdeR \Delta dgcM$ | This work |
| Plasmids |  |  |
| pTrc99a | Expression vector; *P<sub>trc</sub>* promoter inducible by isopropyl-β-D-thiogalactopyranoside (IPTG); pBR ori; Ap<sup>R</sup> | [3] |
| pVS2689 | pTrc99a::*pdeH* | This work |
| pVS1644 | pTrc99a::*dgcE* | This work |

1. Serra DO, Richter AM, Klauck G, Mika F, Hengge R. Microanatomy at cellular resolution and spatial order of physiological differentiation in a bacterial biofilm. *mBio*. 2013;4(2):e00103-13. Epub 2013/03/21. doi: 10.1128/mBio.00103-13. PubMed PMID: 23512962; PubMed Central PMCID: PMC3604763. 2. Besharova O, Suchanek VM, Hartmann R, Drescher K, Sourjik V. Diversification of gene expression during formation of static submerged biofilms by *Escherichia coli*. *Frontiers in Microbiology*. 2016. doi: 10.3389/fmicb.2016.01568. 3. Amann E, Ochs B, Abel K-J. Tightly regulated tac promoter vectors useful for the expression of unfused and fused proteins in *Escherichia coli*. *Gene*. 1988;69(2):301-15. doi: https://doi.org/10.1016/0378-1119(88)90440-4.

## Supporting protocols

### Design and fabrication of the mother machine

A microfluidic device was used for time-resolved studies of *E. coli* growth and curli expression, which features mother machines traps for the observation of one-dimensional cell growth. The design of the microfluidic device is shown in Figure 5. The channel structure of the microfluidic chip features four independent channels with a depth of 8 μm, allowing the realization of four conditions, e.g. different media compositions, in one experiment. Each channel contains two inlets enabling on-chip switching between two different media, which are supplied at defined pressures. By controlling the pressure ratio between the two inlets, the medium supplied at the higher pressure flows through the central channel following the junction, while the other medium is pushed away from the junction and flows out through the waste channel (Figure 5D). The medium which flows through the central channel reaches the mother machine cultivation sites before exiting the chip through a single outlet.

The device was produced using a two-layer soft lithography method as described previously [1]. Based on our in-house made design of the channel layout, a 100 mm silicon wafer was produced by e-beam lithography (ConScience, Sweden). The wafer contains the channel layout as a positive relief. The mother machine traps, which are shown in blue in Figure 5A-C, were structured by etching the wafer by 0.8 μm, giving the mother machine the appropriate vertical dimension for cell trapping. The supply channels, which are shown in green in Figure 5A-C, are implemented as photoresist structures with a height of 8 μm on the wafer. The wafer served as a master mold for liquid polydimethylsiloxane (Sylgard 184 PDMS, VwR International GmbH, Germany), which was mixed at volumetric ratio of 7:1 with a cross-linking agent, degassed in a desiccator for 30 min and poured over the wafer to a height of approximately 4 mm and thermally cured at 80 °C overnight. The cured PDMS was peeled off from the wafer and manually cut into separate chips. Inlet and outlet holes were punched with a 0.75 mm punching tool (Robbins True-Cut Disposable Biopsy Punch 0.75mm with Plunger, Robbins Instruments, USA). The surface of the chip was cleaned by a rinse with isopropanol and the application of adhesive tape (tesafilm, Germany) prior to bonding. The chip was irreversibly bonded to a glass substrate by applying oxygen plasma to both chip and glass surfaces (Diener Femto, Diener GmbH, Germany) and bringing the treated surfaces together. The bond was strengthened by storing the bonded device in the oven at 80 °C for 2 min. The device was mounted on an inverted fluorescence microscope (Nikon Eclipse Ti, Nikon Corporation, Japan) equipped with an incubator. The microscope setup included an Andor Zyla 4.2 sCMOS camera (Oxford Instruments, UK), an objective with 100x magnification (Plan Apochromat λ Oil, NA=1.45, WD=170 μm; Nikon, Japan) and a perfect focus system (Nikon Corporation, Japan) for focus drift compensation.

### Growth experiments

*E. coli* cells were allowed to grow in TB medium until the stationary phase prior to inoculation into the device. Conditioned medium was prepared by cultivating the wild-type *E. coli* cells in TB medium for 20 h (batch culture), after which the cell suspension was centrifuged at 4000 rpm for 10 min. The medium then was filter sterilized and stored at 4°C. The mother machine growth sites were loaded with the undiluted cell suspension by manual infusion of the cell suspension through one of the two inlets using a 1-ml syringe. The connection between the syringe and the chip was realized by tygon tubing (Tygon S-54-HL, inner diameter = 0.51 mm, outer diameter = 1.52 mm, VwR International GmbH, Germany) in combination with blunt dispensing needles (general purpose tips, inner diameter = 0.41 mm, outer diameter = 0.72 mm, Nordson EFD, USA). Medium flow was controlled by programmable pressure regulators (LineUP FlowEZ, FLUIGENT, France), which generated flow by applying pressure on 50-ml medium reservoirs (P-CAP series, FLUIGENT, France). Fresh and conditioned TB medium were respectively filled in separate reservoirs, and each one was pressurized by one module of the pressure regulator. After the cell inoculation both media were connected to the inlets of the channel via tygon tubing and blunt dispensing needles. The pressure at the inlet of the fresh TB medium was set to 200 mbar and remained constant throughout the experiment. During the selection of the positions for imaging the pressure at the inlet of the conditioned medium was set to 250 mbar, allowing the conditioned medium to flow through the junction to the mother machine growth sites and thereby maintaining the stationary state of the cells. At the beginning of imaging, the pressure at the inlet of the conditioned medium was reduced to 150 mbar and programmed to increase back to 250 mbar after 4 h of on-chip cultivation, thereby activating a medium switch from fresh to conditioned TB medium. Phase contrast and GFP fluorescence images were acquired with a time interval of 10 min.

### Analysis of microfluidics data

Cells were segmented from phase contrast images by making use of a fully convolutional neural network based on the U-net architecture [2]. A set of manually curated cell outlines was prepared for training (1105 outlines) and validation (346 outlines) of the network. The training set was augmented by scaling, rotation, flipping and addition of white noise. A U-net of depth three, with 8, 16 and 32 filters along the contracting path, was trained to predict cell interiors from phase contrast images. Phase contrast images were normalized by subtracting the median and scaling to intensities expected between the 2^nd^ and 98^th^ percentiles. Cell interiors were defined from the curated outlines by filling each outline and then subjecting it to two rounds of morphological erosion. The erosion step ensured that neighbouring cells predicted by the network were well separated, such that distinct cell instances could be clearly identified simply by thresholding the prediction and labelling connected regions. After instance identification, two rounds of morphological dilation restored each mask to its original size. Finally, a smooth outline for each cell was obtained as a two-dimensional spline defined by equidistant knots placed on the mask edge.

To track cells between time points, we applied a length conservation strategy for the cells along each trench. At each time point, we ordered cell outlines by their depth in the trench, with deepest cells first. We then attempted to match, in order, a cell outline in time point *t* with one or more cell outlines in time point *t*+1, chosen such that the sum of their cell lengths would be conserved within some threshold tolerance. In the trivial case, the length of the first outline at time point *t* would match that of the first outline at time point *t*+1. In the event of cell division, the cell length at time point *t* would match the sum of the first two cell lengths at time point *t*+1. Since cells may grow in length between time points, we also initialised a growth rate parameter for each cell that biased the expected cell lengths for time point *t*+1 as a fold-increase in length. To enable adaptation to the true growth rate, the growth rate parameter was updated by a lagging average over 20 time points. To increase robustness to errors in segmentation, we additionally allowed state transitions from one to many and many to one, and built a proposal tree, which branched for all valid assignments lying within the length thresholds. We searched for the proposal with the lowest average fold-change in matched lengths, but limited branching by retaining only the 10 best proposals for subsequent nodes (cell outlines) in the tree. The length thresholds were deliberately set loosely such that the (sum of) cell length(s) at *t*+1 could decrease at most five-fold or increase at most two-fold relative to the (sum of) cell length(s) at *t*. This increased the number of valid proposals, but was important in cases where the growth rate estimate was poor. For transitions where one cell outline split into more than two, or transitions where multiple outlines merged into one cell, a new label was generated for the corresponding cells at *t*+1. We made one exception to this labelling strategy to account for occasional ambiguity in segmentation near division events, where a cell segmented as two sister cells could later be segmented as a single mother cell. Specifically, when two sister cells — i.e., cells that were previously involved in a division event — merged into one, the label was set back to that of the mother; at the next division event, the labels of the sister cells were also retained. Finally, note that any outlines below a minimum size threshold of 50 pixels were ignored. All errors in tracking were manually curated.

Cell length was estimated from cell regions as the ‘major axis length’ of the Matlab regionprops function — the major axis of the ellipse with same normalised second central moment as the region. Instantaneous growth rates were estimated from the derivative of a smoothing spline fitted to the logarithm of cell length over each cell division cycle. Knots for the spline were placed at intervals of at most 15 time points. Single-cell fluorescence traces were quantified from the median fluorescence within each outline. Background fluorescence varied as a function of time due to the accumulation of cells at some trench exits, so we corrected for background fluorescence in each trench at each time point using the median value of all non-cell pixels. Fluorescence traces were characterised along branching lineages and were smoothed with a Savitzky-Golay filter of order 3 and window length 21. The derivative of the filter was used to obtain the maximum rate of fluorescence increase. Peaks in fluorescence were identified as points for which the second derivative was less than −100 and either the first derivative was zero, or its sign changed at the next time point. In cases where multiple descendants shared a common peak event before branching, we counted that event only once. The half-life of decay after a peak in fluorescence was determined by fitting an exponential decay function with offset term to fluorescence data after the peak. Given the occasional occurrence of a second peak in fluorescence, the decay function was only fit to time points after the peak up to the point where the first derivative exceeded zero.

